# Nodal root diameter and node number in maize (*Zea mays L.*) interact to influence plant growth under nitrogen stress

**DOI:** 10.1101/2020.12.23.424185

**Authors:** Hannah M. Schneider, Jennifer T. Yang, Kathleen M. Brown, Jonathan P. Lynch

## Abstract

Under nitrogen limitation, plants increase resource allocation to root growth relative to shoot growth. The utility of various root architectural and anatomical phenotypes for nitrogen acquisition are not well understood. Nodal root number and root cross-sectional area were evaluated in maize in field and greenhouse environments. Nodal root number and root cross-sectional area were inversely correlated under both high and low nitrogen conditions. Attenuated emergence of root nodes, as opposed to differences in the number of axial roots per node, was associated with substantially reduced root number. Greater root cross-sectional area was associated with a greater stele area and number of cortical cell files. Genotypes that produced few, thick nodal roots rather than many, thin nodal roots had deeper rooting and better shoot growth in low nitrogen environments. Fewer nodal roots offset the respiratory and nitrogen costs of thicker diameter roots, since total nodal root respiration and nitrogen content was similar for genotypes with many, thin and few, thick nodal roots. We propose that few, thick nodal roots may enable greater capture of deep soil nitrogen and improve plant performance under nitrogen stress. The interaction between an architectural and anatomical trait may be an important strategy for nitrogen acquisition. Understanding trait interactions among different root nodes has important implications in for improving crop nutrient uptake and stress tolerance.

## Introduction

Developing stress tolerant, resource efficient crops is a key strategy for addressing the challenges of climate change, global food security, and land degradation (Blum and Jordan, 1985; Mickelbart *et al.*, 2015; Hunter *et al.*, 2017). Maize is a critical global crop, cultivated for food, fuel, and industrial uses (FAO, 2018). In intensive agriculture systems, nitrogen (N) fertilizers are over-applied to maximize grain yield, yet over half of the applied nitrate leaches beyond the root zone and pollutes waterways, or is volatilized as greenhouse gases (Hirel *et al.*, 2011; Dhital and Raun, 2016). In low-input subsistence agriculture, which sustains half of the global population, maize is grown on marginal soils where nitrogen availability is a primary constraint to yield (Gibbon *et al.*, 2007). Breeding nitrogen-efficient and nitrogen-stress tolerant maize varieties would therefore have substantial economic and environmental benefits.

Root systems have evolved structural and physiological strategies to forage for resources in complex, heterogeneous soil environments (York *et al.*, 2013; Lynch, 2015; Rangarajan *et al.*, 2018). When nitrogen is limiting, plants preferentially allocate resources to root construction and maintenance, as opposed to shoot growth (Brouwer, 1962; Bloom *et al.*, 1985). However, the utility of investing in diverse root strategies is difficult to assess. For example, maize develops spatiotemporally and genetically distinct classes of embryonic and post-embryonic roots, each composed of specialized axial (supportive, conducting) and lateral (branching, absorptive) roots (Hoppe *et al.*, 1986; Demotes-Mainard and Pellerin, 1992; Hochholdinger *et al.*, 2004). Nodal roots emerge acropetally in whorls through development either belowground (“crown”) or aboveground (“brace”) (Hochholdinger et al., 2004). These nodal roots comprise the bulk of the axial root system, and, with their lateral roots, are responsible for the majority of water and nutrient uptake (Schneider *et al.*, 2020). In addition, nodal root position influences size-related traits, like diameter and number, which increase with younger node positions (i.e. nodes of a younger ontogenetic age) (York and Lynch, 2015). Although it is known that roots of a larger diameter are associated with younger nodal positions, there is still substantial genetic variation for root diameter and number of roots at these nodal positions. The understanding and development of a phene (‘phene’ is to ‘phenotype’ as ‘gene’ is to ‘genotype) (York *et al.*, 2013) for crop improvement is challenging and requires understanding of the fitness landscape, which for roots is largely unknown and likely to be highly complex. Understanding the utility of root phenes in variable and dynamic nitrogen regimes, phene tradeoffs for contrasting soil resources, and phene interactions may facilitate the incorporation of root phenotypes in crop breeding (Lynch, 2015, 2018).

Several root ideotypes, or integrated root phenotypes defined as breeding targets, have been proposed for improving nitrogen acquisition efficiency (York *et al.*, 2013); for root system ideotypes, see (Mi *et al.*, 2010; White *et al.*, 2013; Lynch, 2013, 2015, 2018; Lynch and Wojciechowski, 2015; Schmidt and Gaudin, 2017). To understand how different root phenes contribute to nitrogen stress adaptation, whole root system responses to nitrogen stress, as well as the utility of individual phenes, have been explored. Phene states such as steep crown root angle, fewer nodal roots, increased root cortical aerenchyma, and reduced lateral root branching have been associated with rapid, deep rooting and better yield under nitrogen stress among recombinant inbred lines (RILs) (Trachsel *et al.*, 2013; Saengwilai *et al.*, 2014*b*,*a*; Zhan and Lynch, 2015; Lynch and Wojciechowski, 2015), although modeling results suggest that phenes which maximize deep rooting may only provide benefits under precipitation regimes and soil textures that facilitate nitrate leaching (Dathe *et al.*, 2016). Under low nitrogen conditions, some maize genotypes reduced the number and diameter of nodal roots, developed steeper root angles, increased the ratio of lateral to nodal root length, and increased expression of nitrate transporters, among other changes (Gaudin *et al.*, 2011*a*,*b*; Trachsel *et al.*, 2013; Gao *et al.*, 2015). Over time, maize breeding has indirectly resulted in increasing nitrogen use efficiency (Ciampitti and Vyn, 2012; York *et al.*, 2015; DeBruin *et al.*, 2017).

A comparison of commercial maize hybrids over the last century demonstrated that the most recent varieties had multiple changes in root phenotypes, including fewer (per node) but shallower nodal roots, delayed lateral root branching, and increased metaxylem vessel number and smaller diameter vessels. The functional-structural model *SimRoot* demonstrated that a reduction in nodal root number makes century-old genotypes as productive as modern root phenotypes in modern production environments (York *et al.*, 2015). However, the utility of root phenes varies with nutrient regimes and depends on their fitness landscape, including its interactions with other phenes.

Understanding and accounting for phene interactions is important in determining the utility of a phene state for resource capture. Phenes that are beneficial for soil resource acquisition but have a high metabolic cost may have important interactions with root phenes that reduce the metabolic cost of resource capture. The formation of root cortical aerenchyma reduces the metabolic cost of the root and enables more carbon and nitrogen resources to be allocated to root growth (or formation of more nodal roots) for increased capture of nitrogen (Saengwilai *et al.*, 2014*a*,*b*). For example, increased root cortical aerenchyma formation interacts with production of more nodal roots, increasing plant growth in nitrogen limiting conditions up to 130% more than the expected additive effects (York *et al.*, 2013). In common bean, root hair length and basal root growth angle interact in low phosphorus conditions, by affecting the placement of root resources in specific soil domains and increased plant growth up to twice the expected additive effect (Miguel *et al.*, 2015). In common bean, simulations demonstrated that a greater number of basal root whorls and greater lateral root branching density was optimal in low phosphorus environments because more roots developed in the topsoil, which has greater phosphorus availability. In contrast, fewer basal root whorls and decreased lateral root branching enables deeper root growth and better nitrate capture (Rangarajan *et al.*, 2018). A steep growth angle increases nitrogen capture in environments where nitrogen is available deep in the soil profile (Trachsel et al. 2013; Dathe et al. 2016). *In silico*, root hair length, density, geometry, and initiation show substantial synergism for P capture in Arabidopsis (Ma *et al.*, 2001). Phene synergisms show strong interactions with environmental conditions, therefore may be important drivers in evolution and should be considered in crop breeding.

To understand the effects of combining potentially adaptive root traits, we evaluated the relationship of two root phenotypes – the number of nodal roots (a combination of the number of developed root nodes and the number of roots per node), and nodal root diameter (measured as root cross-sectional area). While fewer nodal roots have been shown to improve nitrogen stress tolerance, the utility of anatomical traits, such as root cross-sectional area, within the context of different nodal root number has not been studied. Previous work has suggested a benefit of maintaining larger root cross-sectional area under low nitrogen (Yang *et al.*, 2019), despite increased resource costs. However, interactions of a larger root cross-sectional area with other root traits, such as a simultaneous reduction in nodal root number, could potentially offset resource costs (e.g. nitrogen and carbon) and be beneficial for nitrogen acquisition. For example, anatomical phenes that influence root cross-sectional area are strong predictors of penetration strength of hard soils (Chimungu *et al.*, 2015), root longevity and resilience (Eissenstat *et al.*, 2000), and may enable greater hydraulic conductance. Here we investigate a novel interaction between an architectural trait, fewer nodal roots, and an anatomical trait, a reduced cross-sectional area, for enhanced nitrogen acquisition. We hypothesize that 1) genotypes with attenuated emergence of root nodes produce fewer, thicker nodal roots and 2) fewer nodal roots offset the respiratory and nitrogen costs of thicker diameter roots, resulting in greater plant performance and root depth under nitrogen stress. Maize recombinant inbred lines (RILs) contrasting in both nodal root number and root cross-sectional area were used to evaluate combined trait effects on root respiration, nitrogen content, root length, root depth distributions, and shoot growth, in greenhouse and field experiments under high and low nitrogen conditions.

## Materials and Methods

### Plant material and growth conditions

A greenhouse experiment (8 RILs) and a field experiment (11 RILs) were performed in 2015. Experiments were performed using RILs from the IBM population Maize seeds were provided by Dr. Shawn Kaeppler from the University of Wisconsin. Genotypes from all greenhouse and field studies are listed in Supplementary Table S1.

Greenhouse mesocosm experiments were conducted in the greenhouse at University Park, PA (40° 45’ 36.0” N, 73° 59’ 2.4” W), with 14 h photoperiod, maintained at approximately 28°C/26°C day/night, 40% RH, PPFD of 500 μmol m^−2^ s^−1^ at the sixth leaf (Growmaster Procom, Micro Grow, Temecula, CA, USA). Germination and harvesting dates are listed in Supplementary Table S1. Seeds were surface sterilized with 25% (v/v) commercial bleach for 3 min, rinsed with distilled water, then soaked in the seed fungicide Captan (0.2 g/L) for at least 10 min. Seeds were placed 2.5 cm apart in a row between two sheets of heavy weight seed germination paper (Anchor Paper Co., St. Paul, MN, USA), then rolled up and placed vertically in an imbibing solution of 0.5 mM CaSO_4_, and dark-incubated at 28°C for 2 days. Representative seedlings from each genotype were transplanted at about 5 cm depth.

Plants were grown in four replications in individual mesocosms consisting of polyvinyl chloride cylinders with an inner diameter of 15.5 cm and height of 1.54 m and lined with transparent 6 mm high-density polyethylene film to facilitate root sampling. Genotypes were grown in a two-way factorial complete block design. Each mesocosm was filled with a 30 L mixture consisting of 50% commercial grade medium sand (Quikrete, Harrisburg, PA, USA), 27% horticultural grade fine vermiculite (D3, Whittemore Companies Inc., Lawrence, MA, USA), 18% field soil, and 5% horticultural grade super coarse perlite (Whittemore Companies Inc.), by volume. Soil was collected from the top 20 cm of low-nitrogen fields (Hagerstown silt loam: fine, mixed, semi-active, mesic Typic Hapludalf) maintained at the Russell E. Larson Agricultural Research Center at Rock Springs, PA, air dried, crushed and sieved through a 4 mm mesh. A thin surface layer of perlite was added to each mesocosm to help retain moisture and reduce compression of media.

Plants were fertigated with high (7300 μm) or low (140 μm) N nutrient solutions using drip rings (formulae in Supplementary Table S2). Nutrient solutions were adjusted to pH 6.0 using KOH pellets, and maintained at this pH with KOH or HCl as needed. A dilute micronutrient foliar spray was applied uniformly as needed. Each mesocosm was saturated with 2.5 L of nutrient solution one day prior to transplant, then fertigated 200 mL per mesocosm every other day.

The field trials were conducted in 0.4 ha fields maintained with split high and low nitrogen treatments at The Pennsylvania State University’s Russell Larson Research Farm (40°42’40.915”N, 77°, 57’11.120”W, which has Hagerstown silt loam soil (fine, mixed, semi-active, mesic Typic Hapludalf). To generate low N conditions, approximately 84 metric tons per ha sawdust was tilled into the soil in 2012 and 2013. High nitrogen fields were fertilized with 157 kg N ha^−1^ urea (46-0-0). The low nitrogen treatment did not receive any nitrogen fertilizer. Seeds were planted in rows with 76 cm row spacing at a density of approximately 56,800 plants ha^−1^. Genotypes were planted in 3-row plots in four replications per genotype in a split-plot randomized block design with 88 total plots. Plants were sampled from the middle row of each plot. Planting and sampling dates are listed in Supplementary Table S1. The average percent reduction due to nitrogen stress in shoot biomass and yield from all experiments is listed in Supplementary Table S3. The soil nitrate distribution by depth under high and low nitrogen treatments is shown in Supplementary Fig. S1.

### Plant sampling and measurements

In greenhouse studies, shoots were removed, dried at approximately 70 °C for 72 h, and stem and leaves were weighed separately. Whole root systems in media were removed intact within polyethylene liner bags and media was gently washed off with a hose. Nodal roots were counted, and lengths measured by node. Two representative root segments each were excised from the second, third, and fourth nodes and preserved in 75% ethanol for anatomical analysis. Root respiration rate was measured on three 2-cm nodal root segments (4 to 6 cm from the stem base) from each of these nodes, using a LI-COR 6400XT (LI-COR Biosciences, Lincoln, NE, USA) fitted with a closed custom chamber within three minutes after harvest. A subset of these roots was dried at about 70°C for 72 h, weighed to determine specific root length, manually ground, and a 2 mg subsample was analyzed for carbon and nitrogen content using a CHN elemental analyzer (2400 CHNS/O Series II, PerkinElmer, Waltham, MA, USA). Each 30 cm of the entire root system, beginning at the stem base, was collected and dried at approximately 70 °C for 72 h to obtain total root biomass. Dried leaves were ground and a 2 mg subsample was analyzed for total nitrogen content as above.

In field studies, three representative plants from each plot were excavated using a shovel (Trachsel et al., 2011). Root crowns were separated from the shoots, soaked in water with 0.5% v/v detergent (Liquinox, Alconox, Inc. USA) for approximately 10 minutes, and hosed to remove remaining soil. Each node of roots was excised, node one being defined as the coleoptilar node, and 3 representative roots from nodes 2, 3, and 4 in the greenhouse and nodes 1, 2, and 3 in the field were sampled at 2 to 4 cm from the base of the stem. Nodes were identified by dissecting from the youngest nodes to the oldest nodes. Anatomical samples were preserved in 75% ethanol for anatomical processing.

For a subset of plants, nodal roots (>2 cm) were counted by node and recorded as crown (emerging below the soil line and having no pigmentation) or brace roots (emerging above the soil line and having pigmentation). Shoot biomass was separated into stem, leaves, and ears, dried at approximately 70°C for 72 h, and weighed. Dried leaves were ground and analyzed for total nitrogen content with a CHN elemental analyzer. Ears from five plants per plot were collected at physiological maturity, dried to approximately 15% moisture content, shelled and weighed.

To determine relative root lengths and depths, soil cores 60 cm in depth and 5 cm in diameter were taken manually with a sledgehammer using a steel coring tube and plastic liner (Giddings Machine Co., Windsor, CO, USA), between two plants in each plot. Soil cores were separated into 10 cm segments and roots were extracted using a custom root washer. Roots were placed in water on a clear plastic tray, scanned (Epson Perfection V700 Photo, Epson America, Inc., Long Beach, CA, USA) at 600 dpi and analyzed for root length separated by diameter classes (e.g. 0.2, 0.5, 1 mm) to estimate lateral versus nodal (i.e. axial) root lengths using Winrhizo Pro software (Regent Instruments, Québec City, Quebec, Canada).

In greenhouse experiments, media samples were collected at different depths in the mesocosm, air-dried, homogenized, and samples of equal mass were tested for nitrate content was using a LAQUA nitrate meter (Spectrum Technologies, Aurora, IL, USA) according to manufacturer’s instructions. For field experiments, soil cores were taken, separated by depth, homogenized, dried at 70°C for 72 h, and extracted as above. The low nitrogen treatment had on average 36% reduced soil nitrogen in the greenhouse and 66% reduced soil nitrogen in the field compared to the high nitrogen treatment (Supplementary Fig. S1).

The nitrogen treatment was confirmed in all experiments by a ~46% vegetative biomass growth reduction in greenhouse experiments, a ~30% vegetative biomass growth reduction in field experiments, and a ~20% yield reduction in field experiments in nitrogen stress compared to control conditions (Supplementary Table S3).

### Image and statistical analysis

For field experiments, the middle portion of two representative root segments per node of each plant were ablated and imaged using laser ablation tomography (LAT) (Hall *et al.*, 2019; Strock *et al.*, 2019). In brief, the root is moved into the laser beam of a nanosecond pulsed UV laser (Avia 355-7000, Coherent, Santa Clara, CA, USA) at about 30 μm s^−1^ and as each surface is ablated and exposed, images are captured. Image scale was 1.173 pixels per micron. Select greenhouse-grown root segments required pre-processing in a critical point dryer (Leica EM CPD300, Leica Microsystems, Wetzlar, Germany) to prevent sample desiccation during laser ablation. Roots were placed in histo prep tissue capsules (Fisherbrand, Fisher Scientific, Waltham, MA, USA) and gradually dehydrated 75% to 100% ethanol prior to critical point drying.

For greenhouse experiments, two ethanol-preserved roots from each node (2, 3, and 4) were manually sectioned using fresh double-edged razor blades, wet-mounted and visualized using a Diaphot inverted light microscope (Nikon Inc., Melville, NY, USA) under 4X magnification with a mounted CCD camera (Nikon DS-Fi1 camera with DS-U2 USB controller, Nikon, Inc.). Images were captured using NIS Elements F 4.30.00 software (Nikon, Inc.) at a scale of 390.7 pixels per mm, using 1280 x 920 pixel resolution. Two representative cross-sections images per root were selected for analysis.

Images were analyzed using custom macros created with the open-source ObjectJ plug-in in ImageJ (Rasband, 2015) in which cortex, stele, aerenchyma, vessel and cell outlines were manually traced, and cell files manually counted (Supplementary Fig. S2). This allowed careful quantification of cell and vessel sizes. A total of 17 anatomical phenes were measured on each root cross-sectional image (Supplementary Fig. S2).

Statistical analysis and visualizations were generated using R version 3.3.1(R Core Team, 2018). Data were analyzed by linear regression, Pearson correlation coefficients, and Tukey's HSD (honest significant difference). Few, thick phenotypes are classified as having a root cross-sectional area greater than 2 µm and a nodal root number less than 15. Many, thin phenotypes are classified as having a root cross-sectional area less than1 μm and a nodal root number greater than 30. Phenotypes not meeting these criteria were excluded from these analyses. Bar plots were generated using data aggregation functions from the package *plyr* (1.8.6) and plotting functions from the package *ggplot2* (3.0.0) (Wickham, 2016).

## Results

### Reduced nodal root number was associated with increased cross-sectional area among maize RILs

Genotypes with a greater nodal root cross-sectional area, averaged from nodal roots in the second and third nodes, had a reduced number of nodal roots among RILs in high and low nitrogen conditions in the greenhouse and field (Fig. 1 AB). Differences in nodal root number were driven by the number of root nodes, rather than nodal occupancy (Supplementary Fig. S3). Contrasts in root cross-sectional area among maize RILs were most strongly related to cortical cell file number and stele area, rather than cortical cell diameter (Fig. 2). Cortical cell diameter was strongly positively correlated to root cross-sectional area under low nitrogen, but not high nitrogen conditions (Fig. 2A).

**Figure 1.**
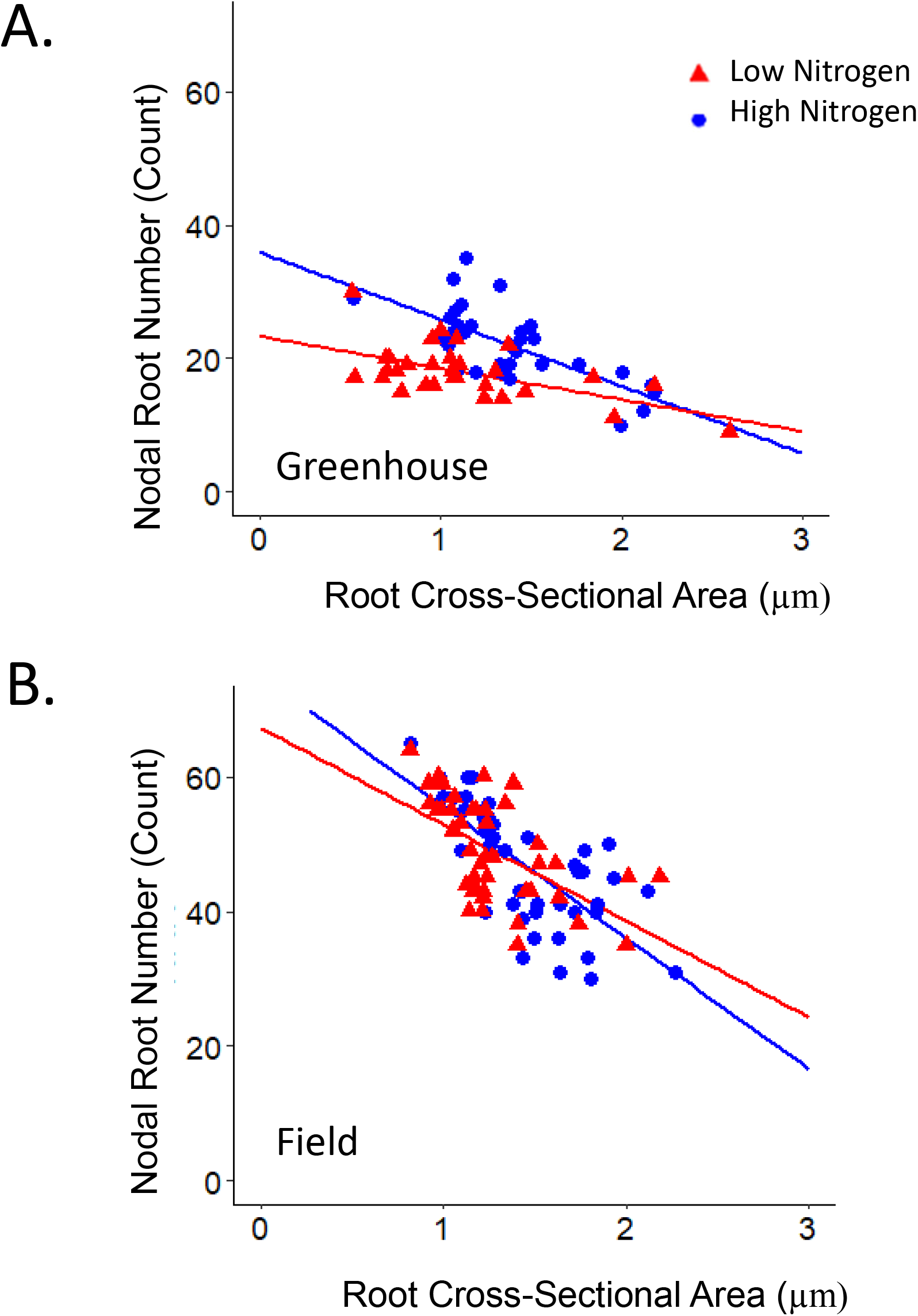
Relationship between nodal root number and root cross-sectional area among maize RILs. Linear regression of number of nodal roots emerged at sampling and root cross-sectional area averaged rom two second and third node roots, from individual plants of IBM RILs grown in high (HN, blue) or ow nitrogen (LN, red) treatments in the: (A) greenhouse (HN: R^2^=0.48, p=0.003; LN: R^2^=0.3, p= 0.004) and (B) field (HN: R^2^=0.5, p=0.003; LN: R^2^=0.32, p= 0.002).

**Figure 2.**
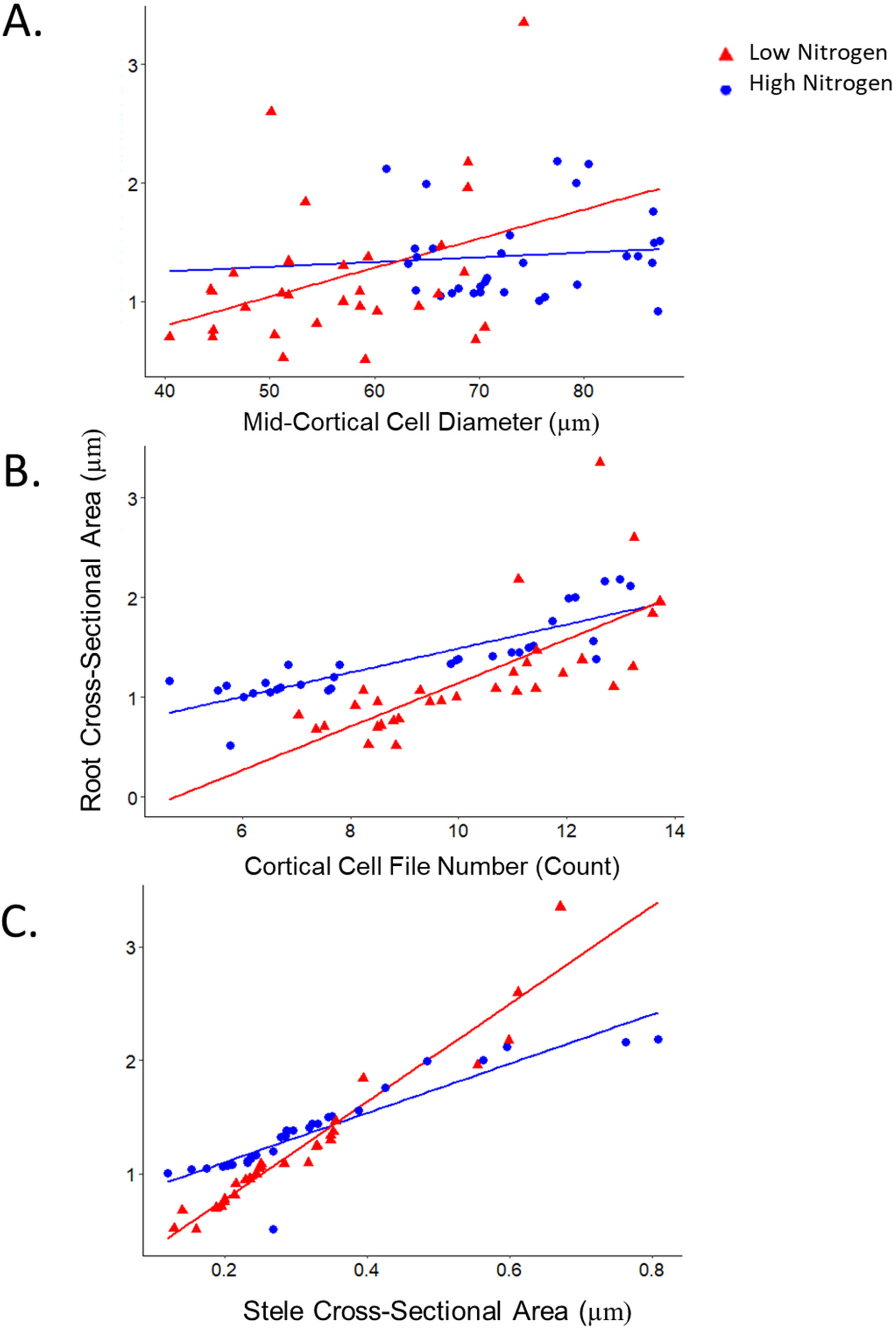
Relationship between nodal anatomical traits and root cross-sectional area among maize IBM RILs. Linear regression of the following nodal anatomical traits against root cross-sectional area averaged from two second and third node roots (fully developed in all genotypes at sampling), from individual plants of maize IBM RILs grown in high HN blue) or low nitrogen (LN, red) treatments in the greenhouse: (A) mid-cortical cell diameter (HN: R^2^=0.001, p=0.004; LN: R^2^=0.13, p= 0.001), (B) cortical cell file number (HN: R^2^=0.72, p=0.004; LN: R^2^=0.5, p= 0.003), and C) stele cross-sectional area (HN: R^2^=0.081, p=0.003; LN: R^2^=0.93, p= 0.002).

### Fewer, thicker nodal roots were associated with better shoot growth under nitrogen stress in maize RILs

Fewer nodal roots were correlated with greater shoot mass under nitrogen stress (Fig. 3A). In contrast, there was no significant relationship between nodal root number and dry shoot biomass under high nitrogen in these studies (Fig. 3A). Larger root cross-sectional area was positively correlated with shoot biomass under low and high nitrogen conditions (Fig. 3B). Lines with the few, thick nodal root phenotype had significantly greater dry shoot biomass in low nitrogen conditions when compared to lines with the many, thin nodal root phenotype (Fig. 4A).

**Figure 3.**
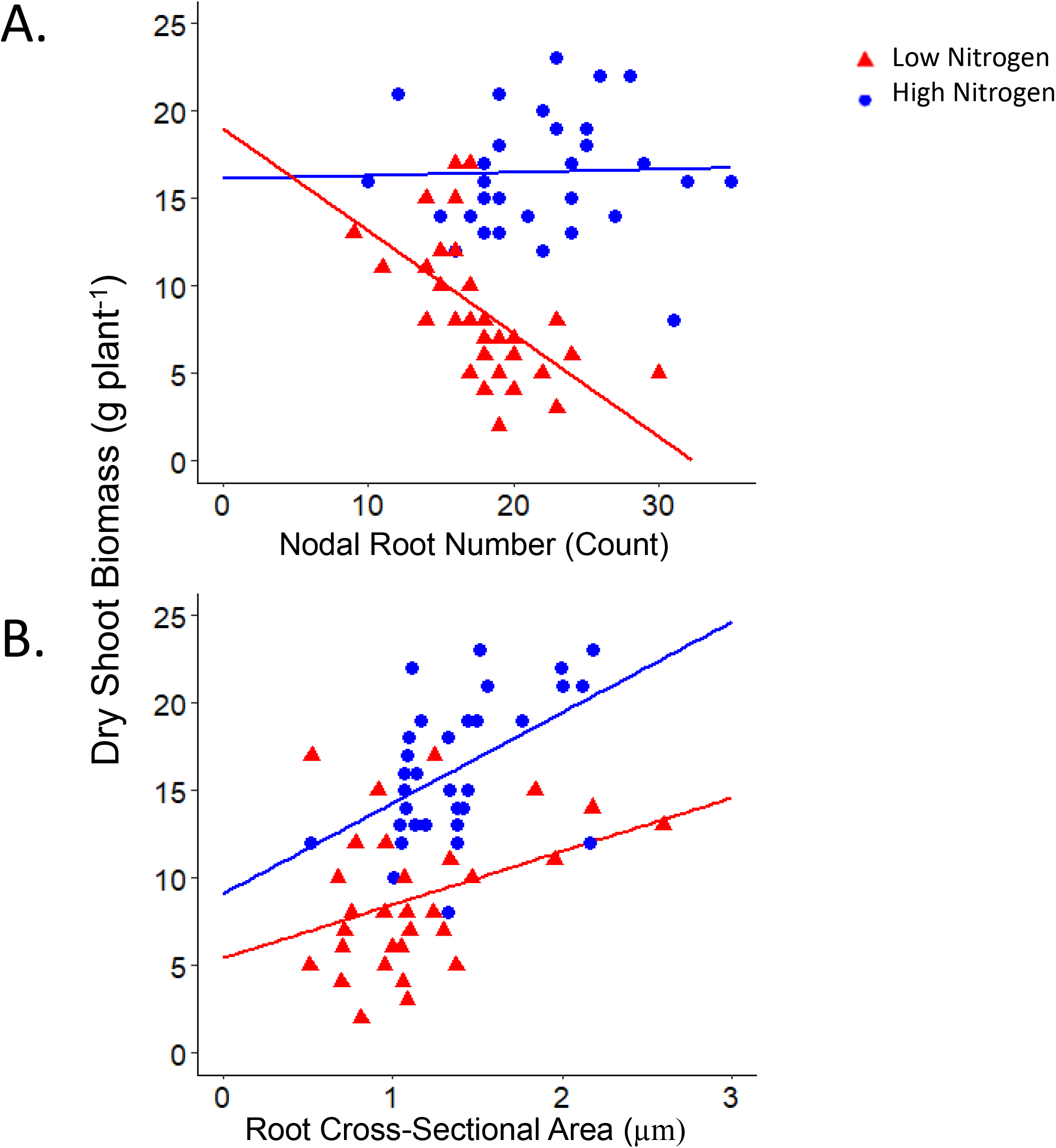
Relationship between shoot biomass, nodal root number and root cross-sectional area in maize IBM RILs. Linear regression of (A) total number of nodal roots at sampling (HN: R^2^=0.11, p=NS; LN: R^2^=0.35, p= 0.003) and (B) root cross-sectional area (HN: R^2^=0.25, p=0.003; LN: R^2^=0.19, p= 0.002) averaged from two second and third node roots, against total dry shoot biomass, from individual plants of maize IBM RILs grown in high (HN, blue) or moderate low nitrogen (LN, red) treatments in the greenhouse

**Figure 4.**
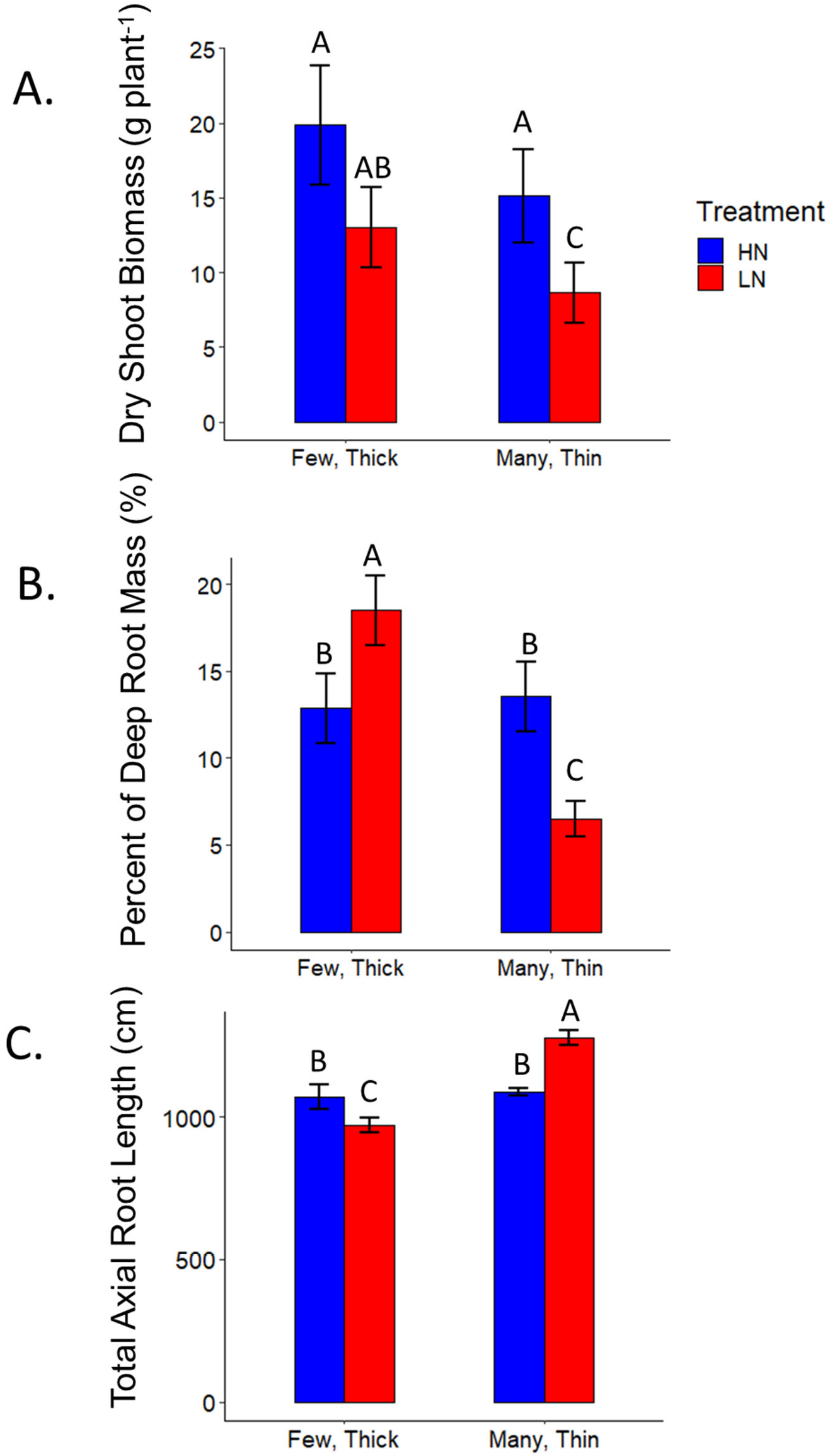
Relationship between shoot biomass, total axial root length, deep root mass, and root phenotypes among maize RILs. A) Lines with few, thick nodal roots have greater dry shoot biomass in low nitrogen compared to lines with many, thin nodal roots. B) Lines with few, thick nodal roots have greater deep root mass in low nitrogen conditions compared to lines with many, thin nodal roots. C) Lines with few, thick nodal roots have a reduced total axial root length when compared to lines with many, thin nodal roots. Few, thick phenotypes are classified as having a root cross-sectional area greater than 2 μm and a nodal root number less than 15. Many, thin phenotypes are classified as having a root cross-sectional area less than1 μm and a nodal root number greater than 30. Phenotypes not meeting this criteria were excluded from these analyses. Data shown are means ± standard error (SE) for three genotypes per group averaged from second and third nodal roots (n=24). Means with the same letters are not significantly different (p ≤ 0.05) according to Tukey’s HSD).

### Genotypes with fewer, thicker nodal roots produced less total nodal root length but deeper roots

The total axial root length produced (the product of nodal number and the average root length) was most strongly related to nodal root number under high and low nitrogen conditions, whereas root cross-sectional area was only weakly negatively correlated with total root length under low nitrogen (Fig. 5AB). Lines with the few, thick phenotype had significantly reduced total axial root length in low nitrogen conditions when compared to lines with many, thin nodal roots (Fig. 4C). Larger root cross-sectional area and fewer nodal roots were significantly correlated with deeper relative root distribution among RILs in low nitrogen conditions (Fig. 4B, 5CD). Lines with the few, thick nodal root phenotype had significantly greater deep root mass in low nitrogen conditions when compared to lines with the many, thin nodal root phenotype (Fig. 4B). Under nitrogen stress, a greater proportion of the root system became deeply distributed. The percent of roots in the shallowest 20 cm decreased, while the percent of roots in the deepest layers increased (Fig. 6).

**Figure 5.**
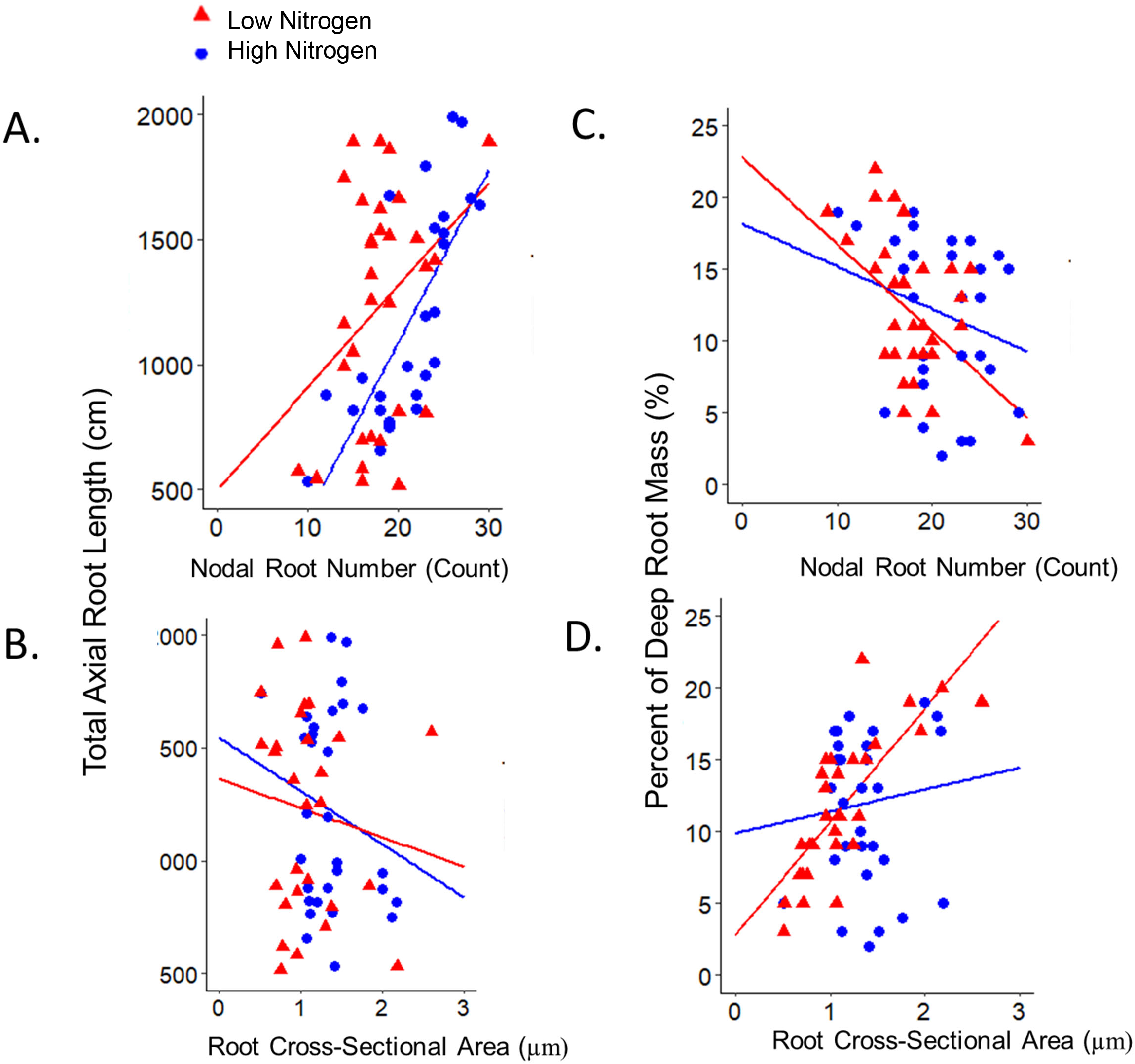
Relationship between total axial root length, deep root mass, root number and cross-sectional area among maize RILs. Linear regression of (A) total number of nodal roots sampled (HN: R^2^=0.03, p=NS; LN: R^2^=0.23, p= 0.003) at harvest against the percent of total root biomass below 124 cm and (B) root cross-sectional area sampled (HN: R^2^=0.013, p=NS; LN: R^2^=0.57, p= 0.004) averaged from two second and third node roots and (C) total number of nodal roots sampled (HN: R^2^=0.05, p=NS; LN: R^2^=0.06, p= 0.005) at harvest and (D) root cross-sectional area sampled (HN: R^2^=0.05, p=NS; LN: R^2^=0.07, p= 0.005) averaged from two second and third node roots, against total axial root length summed from node 2 through all developed root nodes, from individual plants of eight maize IBM RILs grown in high (HN, blue) or moderate low nitrogen (LN, red) treatments in the greenhouse. Plants with missing values were excluded.

**Fig. 6.**
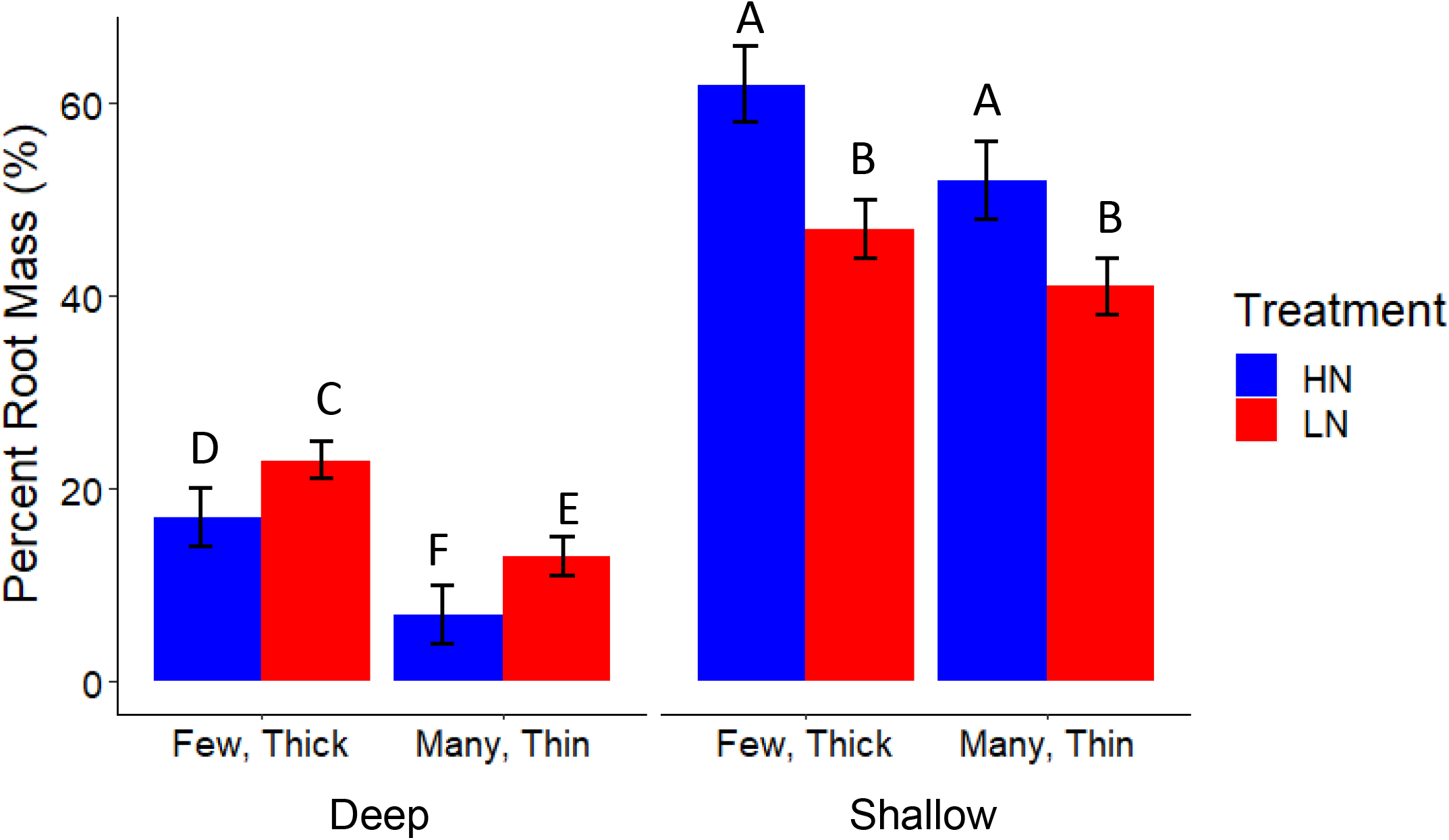
Relative root depth distributions among maize IBM RILs. Means ± SE of the percent of total root mass located below 124 cm (deep) percent of total root mass in the top 30 cm of media (shallow) among IBM RILs in the greenhouse. High and low nitrogen treatments are indicated (HN, blue; LN, red). Few, thick phenotypes are classified as having a root cross-sectional area greater than 2 μm and a nodal root number less than 15. Many, thin phenotypes are classified as having a root cross-sectional area less than1 μm and a nodal root number greater than 30. Phenotypes not meeting this criteria were excluded from these analyses. Means with the same letters are not significantly different (p ≤ 0.05) according to Tukey’s HSD).

### Nitrogen stress effects on root system phenotypes were node-specific

In greenhouse grown plants, nitrogen stress reduced average nodal root length, root cross-sectional area, and nodal occupancy overall, with effects differing by node (Supplementary Fig. S4ABC). In high nitrogen conditions, axial root length decreased in each node; roots from the first two nodes typically reached the bottom of the mesocosm (150 cm) by sampling, five weeks after germination (Supplementary Fig. S4A). Root cross-sectional area increased with each younger root node across genotypes, and nodal occupancy increased from the third node onward. In low nitrogen, fewer plants, particularly plants that develop fewer nodal roots, had developed a sixth node resulting in an apparent increase in nodal occupancy in node six (2 plants measured in high nitrogen, 1 plant measured in low nitrogen) (Supplementary Fig. S4BC).

### Fewer nodal roots offset increased carbon and nitrogen costs of thicker nodal roots

Nitrogen stress significantly decreased nodal root respiration per unit of root length, and in combination with reductions in nodal root length, resulted in substantial reductions in total nodal root respiration per plant (Fig. 7 AB). Similarly, reduced root nitrogen content combined with decreased root biomass resulted in substantial reduction in the percent root nitrogen and total root nitrogen (grams per plant) across genotypes (Fig. 7 CD). Root respiration per unit of root length was significantly positively related to root cross-sectional area in high and low nitrogen conditions (Supplementary Fig. S5). However, when multiplied by the total number and length of nodal roots, total nodal root respiration was similar for genotypes with many, thin and few, thick nodal roots (Figure 7AB). Total root nitrogen content also did not significantly differ between phenotypes when multiplied by root mass indicating that fewer nodal roots offset increased carbon and nitrogen costs of thicker nodal roots (Fig. 7BD). Total root nitrogen was similar across genotypes under nitrogen stress (Fig. 7D).

**Fig 7.**
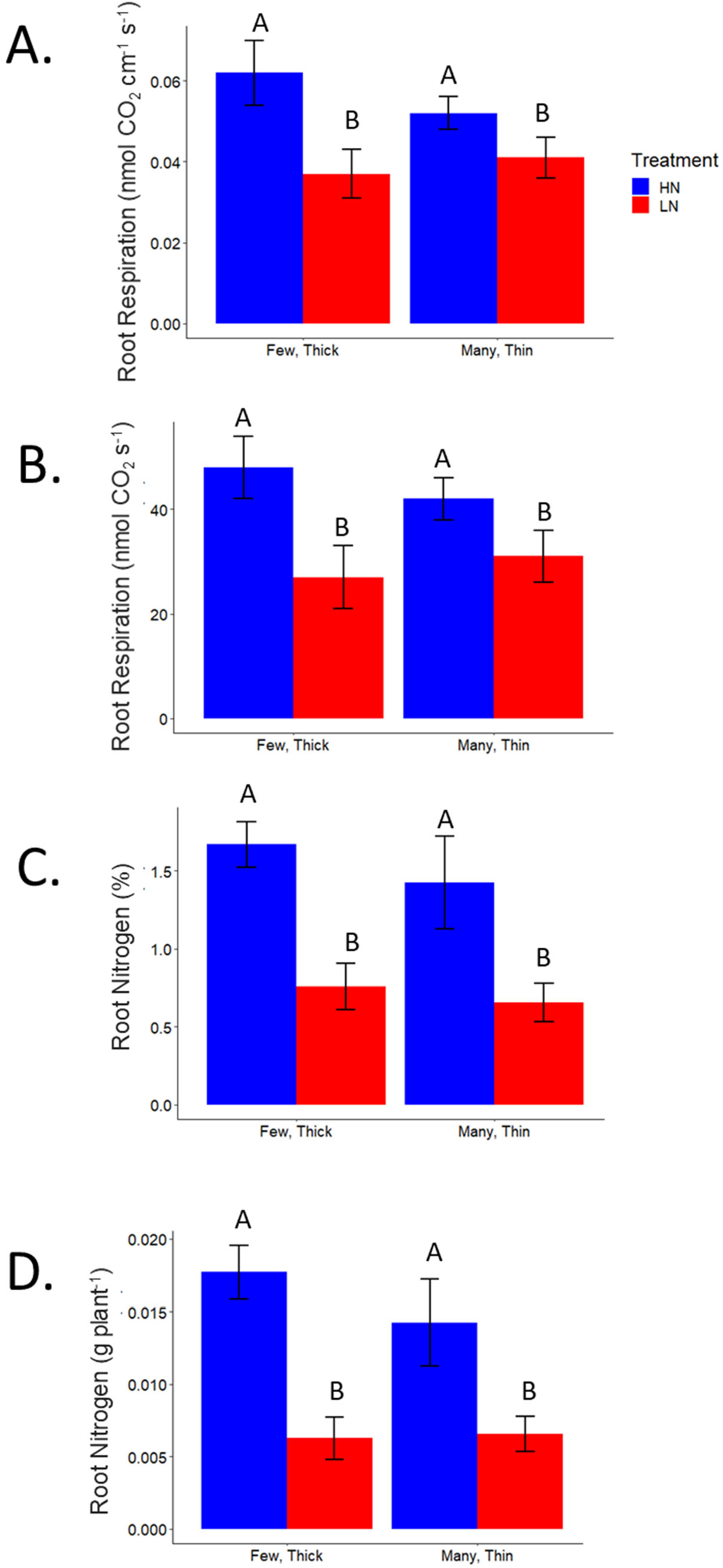
Root respiration and nitrogen content among IBM RILs. Means ± SE of (A) root respiration rates, averaged from six axial roots from each genotype, 2 to 5 cm from stem base, from nodes 2 and 3, and (B) total root respiration (estimated as root respiration rate multiplied by total axial root length in the given node) of nodes two and three, in the greenhouse. Means ± SE of (C) root nitrogen content (percent by mass) from the top 30 cm of the root system (all root classes, homogenized), evaluated in a subset of replicates, and (D) total root nitrogen in the top 30 cm of the root system (percent root nitrogen multiplied by dry root mass in the top 30 cm), in the greenhouse. High and low nitrogen are indicated (HN, blue; LN, red). Few, thick phenotypes are classified as having a root cross-sectional area greater than 2 μm and a nodal root number less than 15. Many, thin phenotypes are classified as having a root cross-sectional area less than1 μm and a nodal root number greater than 30. Phenotypes not meeting this criteria were excluded from these analyses. Means with the same letters are not significantly different (p ≤ 0.05) according to Tukey’s HSD).

## Discussion

This study explored the interaction of root anatomical and architectural traits for nitrogen capture in maize, and identified phenotypes associated with improved plant performance in nitrogen stress: a reduced number of developed root nodes (and thus total number of nodal roots emerged in a given period) and increased nodal root cross-sectional area. Previously, the utility of reduced nodal root number was demonstrated to have utility for enhanced nitrogen capture (Saengwilai *et al.*, 2014*b*). However, here we identified a previously uncharacterized interaction between nodal root number and cross-sectional area to further improve nitrogen acquisition and nitrogen stress tolerance. Genotypic contrasts in root cross-sectional area were driven primarily by differences in cortical cell file number and stele area, with some differences in cortical cell diameter under low nitrogen (Fig. 2). Genotypes with fewer, thicker roots developed root nodes more slowly, produced less total nodal root length, and invested more carbon and nitrogen per unit of root length (Fig. 4, 5, 6). The phenotype of fewer, thicker nodal roots was associated with deeper root distribution and resulted in greater shoot growth under nitrogen stress.

The number of crown roots is an important determinant of soil resource capture (Lynch, 2013) and large genetic variation is present for this trait in maize (Gaudin *et al.*, 2011*a*; Trachsel *et al.*, 2011; Saengwilai *et al.*, 2014*b*; Gao and Lynch, 2016; Sun *et al.*, 2018; Guo *et al.*, 2019). Among field-and greenhouse-grown lines, there was a significant, negative correlation between nodal root number and shoot mass under nitrogen stress (average of 50% biomass reduction) (Fig. 3). Fewer nodal roots is advantageous for the capture of mobile nutrients like nitrogen (Saengwilai *et al.*, 2014*b*), and water (Gao and Lynch, 2016), while a larger number of crown roots is advantageous for the capture of phosphorus (Sun *et al.*, 2018) and presumably other immobile nutrients (Lynch, 2019). In low nitrogen conditions, fewer nodal roots is advantageous as more resources will be available for the development of longer, deeper roots resulting in greater nitrogen acquisition, growth and yield when compared to genotypes with many crown roots. Functional-structural modeling in *SimRoot* suggested that the effects of reduced nodal root number on nitrate uptake were similar regardless of whether this reduction came from delaying the emergence of roots (reduced time with a given number of roots), or producing fewer roots per node (York, 2014).

We observed a consistent positive correlation between root cross-sectional area and shoot mass under high and low nitrogen, among greenhouse-grown and field-grown RILs, suggesting a potential benefit of thicker nodal roots (or, a benefit of thicker roots given concurrent decrease in root number, or other unknown linked traits) for nitrogen stress tolerance. Roots with a smaller root cross-sectional area would be expected to have enhanced performance due to a reduced metabolic cost. However, among RILs, fewer nodal roots reduced root carbon and nitrogen costs, offsetting the increased respiratory costs and nitrogen content of thicker root diameter. We propose that fewer, thicker nodal roots have utility in nitrogen stress. Fewer nodal roots offset resource costs of thicker roots while enabling greater hydraulic conductance and soil penetration strength which have utility for nitrogen capture.

Thicker root diameter has been associated with better performance in hybrids in nitrogen stress, partly due to a positive allometric relationship with plant size (see Yang *et al.*, 2019)). Increased Root cross-sectional area has been associated with increased root penetration strength (Striker *et al.*, 2007; Chimungu *et al.*, 2015), increased hydraulic conductance (Jordan *et al.*, 1993), and lodging resistance (Stamp and Kiel, 1992). Larger mid-cortical cell diameters and fewer cortical cell files were associated with reduced metabolic costs per root length, deeper rooting, and enhanced drought tolerance in IBM RILs (Chimungu *et al.*, 2014*a*,*b*).

Seminal, primary, and lateral root classes were not evaluated in detail in this study. Decreases in lateral to nodal root length ratio have been suggested as a potential adaptation to nitrogen stress (Zhan and Lynch, 2015).. Fewer lateral branches decreases intra- and inter-root competition and decreases root respiration costs enabling deeper root growth and nitrogen capture (Zhan and Lynch, 2015). Additional research is needed to reveal whether plants with few, thick roots invested the additional resources in other root classes, such as additional lateral root proliferation, which could result in greater specific root length overall and increased capacity for nitrate uptake.

Nitrogen stress has large effects on root phenotypes. In low nitrogen, root growth angles became steeper (Trachsel *et al.*, 2013), the proportion of root cortical aerenchyma increases (Gao *et al.*, 2015), nodal root diameter increases (Yang *et al.*, 2019), and in this study, total nodal root length is reduced (Fig. 4) and is the number of nodal roots per node (Supplementary Fig. S3). As a result of changes in root number, diameter, angle and length, in the current study, root systems became more deeply distributed under nitrogen stress in the field and greenhouse, in accord with previous studies (Gaudin *et al.*, 2011*a*; Trachsel *et al.*, 2013; Saengwilai *et al.*, 2014*b*; Postma *et al.*, 2014; Zhan and Lynch, 2015; Dathe *et al.*, 2016). The extent to which the nitrogen regime affected root depth distribution differed among genotypes, but all genotypes showed a substantial reduction in the proportion of root mass in the top 30 cm under low nitrogen in the greenhouse. Similarly, nitrogen stress induced a significant reduction in the percent of root length in the top 20 cm of soil in the field, and an increase in the percent of root length in the deepest 20 cm, among RILs. Finally, nitrogen stress induced a well-established increase in root to shoot mass allocation, which can be observed generally as a shift in allometric scaling between root and shoot mass across genotypes. Shoot mass was also weakly correlated to root cross-sectional area under high nitrogen in some experiments, suggesting that a positive allometric relationship of root traits with plant size may exist. The integration of root and shoot responses to nitrogen stress in maize, and the impacts of different anatomical, morphological, and architectural strategies on nitrogen uptake and utilization efficiency, are complex and merit further research.

Nitrogen stress significantly decreased the number of roots per node in all nodes except the first two. However, genotypes varied in the node at which nodal occupancy began to decrease, and some genotypes (e.g. IBM181) showed no significant decrease in occupancy across nodes (Supplementary Fig. S3). Previous studies have shown that timing of the initiation of root node primordia is staggered, and all initiated primordia always elongate in the first five nodes (Sharman, 1942; Girardin *et al.*, 1986; Aguirrezabal *et al.*, 1993). Aguirrezabal and colleagues (1993) also suggested that in contrast to the first five nodes, subsequent nodes regularly initiated excess root primordia which did not elongate, increasing their sensitivity to carbon availability and the potential for plastic responses (Aguirrezabal *et al.*, 1993). If root initiation preceded onset of stress signaling, nodal occupancy would likely not be affected (e.g. Pellerin, 1994) (in contrast to elongation rate, which could therefore be considered more “plastic”). Therefore, the timing and level of nitrogen stress could directly impact the potential for decreased nodal root number..

Our results demonstrate that few, thick nodal roots are beneficial for plant growth when compared to plants with many, thin nodal roots. Small differences in data trends between the field and greenhouse could be attributed to differences in the growth environment, the spatiotemporal location of nitrogen in the soil, and differences in soil bulk density. In both the greenhouse and field, we employed ‘near isophenic’ (i.e. similar in all root traits except for crown root number and root cross-sectional area) RILs to explore the physiological utility of root traits in nitrogen stress. RILs share a common genetic background (i.e. descending from the same two parents) and are suited to the physiological analysis of phenotypes controlled by multiple alleles. RILs are suitable for this study because they are closely related genotypes therefore minimizing the risk of effects from epistasis, genetic interactions, and pleiotropy which may cofound interpretation of results from other related lines (Zhu *et al.*, 2006).

We highlight an integrated root phenotype related to a developmental process (the rate of root node development) which may underlie phenotypic contrasts and physiology. The integration of nodal root number and root anatomy may have important implications for soil resource capture. Root diameter is a strong predictor of penetration strength in hard soils (Chimungu *et al.*, 2015), root longevity and resilience (Eissenstat *et al.*, 2000), and may enable greater hydraulic conductance. In addition, fewer nodal roots enable deeper rooting and potentially reducing the spatial overlap among axial roots and therefore reducing intra-plant competition for soil resources. Understanding trait interactions among different root nodes has important implications in ideotype breeding of crop stress tolerance.

## Abbreviations

IBM: intermated B73 x Mo17
N: nitrogen
RIL: recombinant inbred line

## Acknowledgments

We thank Shawn Kaeppler and the G2F consortium for providing plant materials. For agronomic and technical support, we thank Qun Wang, Robert Snyder, Michael Williams, Johan Prinsloo, Ainiz Fauzi, Jonathan Wu, Pufan Wang, Ryan Burnett, Tania Galindo-Castañeda, Stephanie Klein, Claire Lorts, Gustavo da Silveira, Andy Evensen, Xiyu Yang, Kirsten Lloyd, Larry York, and Joseph Chimungu. We thank Benjamin Hall and support from the Edmund Optics Higher Education Grant for technical support and improvements to the LAT platform. This work was supported by USDA NIFA’s AFRI grant 2013-02682 to JPL, the Howard G. Buffett Foundation for research in South Africa, National Institute of Food and Agriculture, U.S. Department of Agriculture, award #2014-67013-21572, USDOE ARPA-E ROOTS Award Number DE-AR0000821 and Hatch project #4582.

---

**Fig S1.**
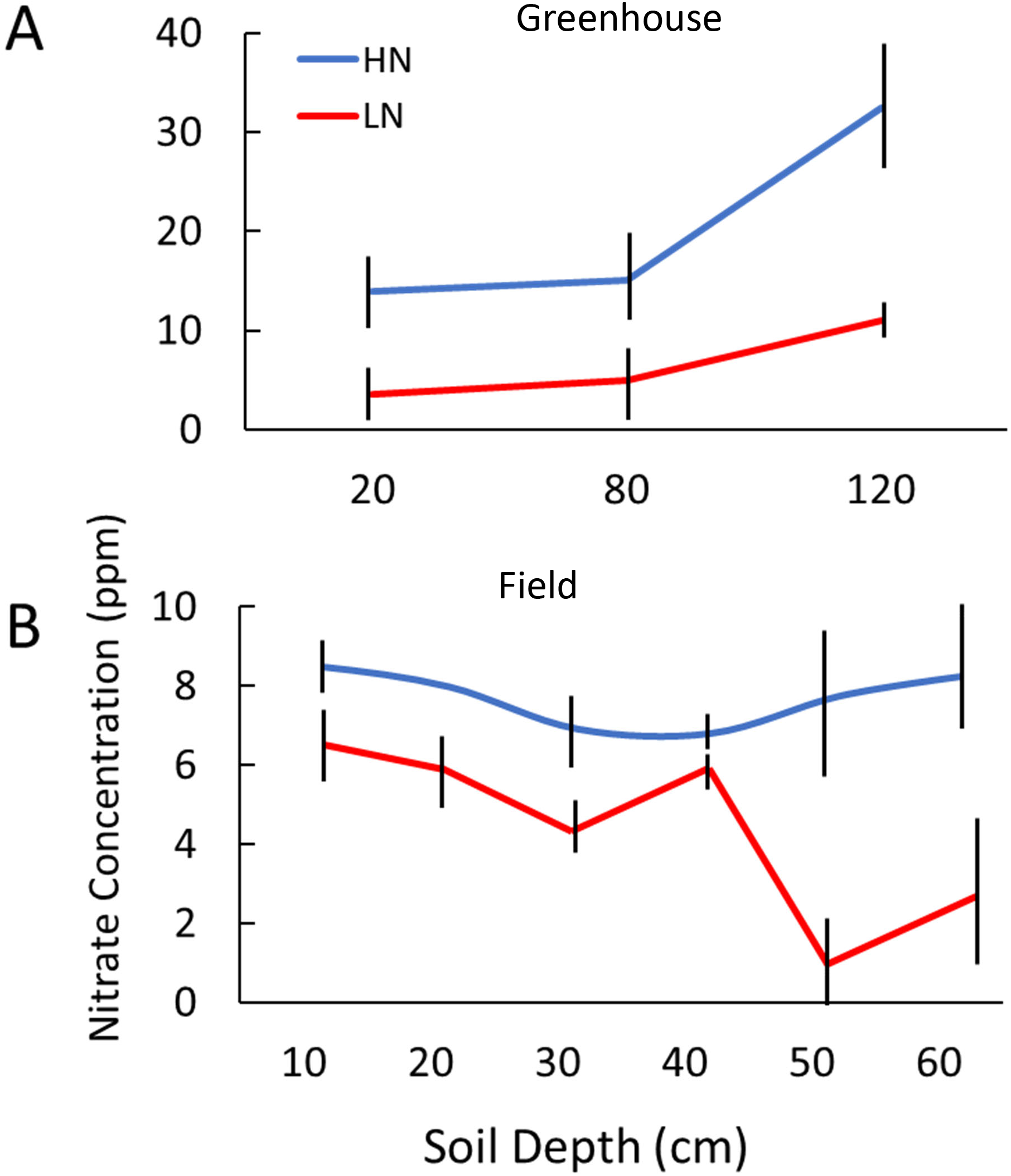
Nitrate concentrations by depth in greenhouse and field experiments. Nitrate concentration from soil or media extracts, averaged across soil or media samples from at least two replicates in the greenhouse and field from the indicated depths in high (HN, blue) or low (LN, red) nitrogen treatments. Error bars represent the standard error.

**Fig S2.**
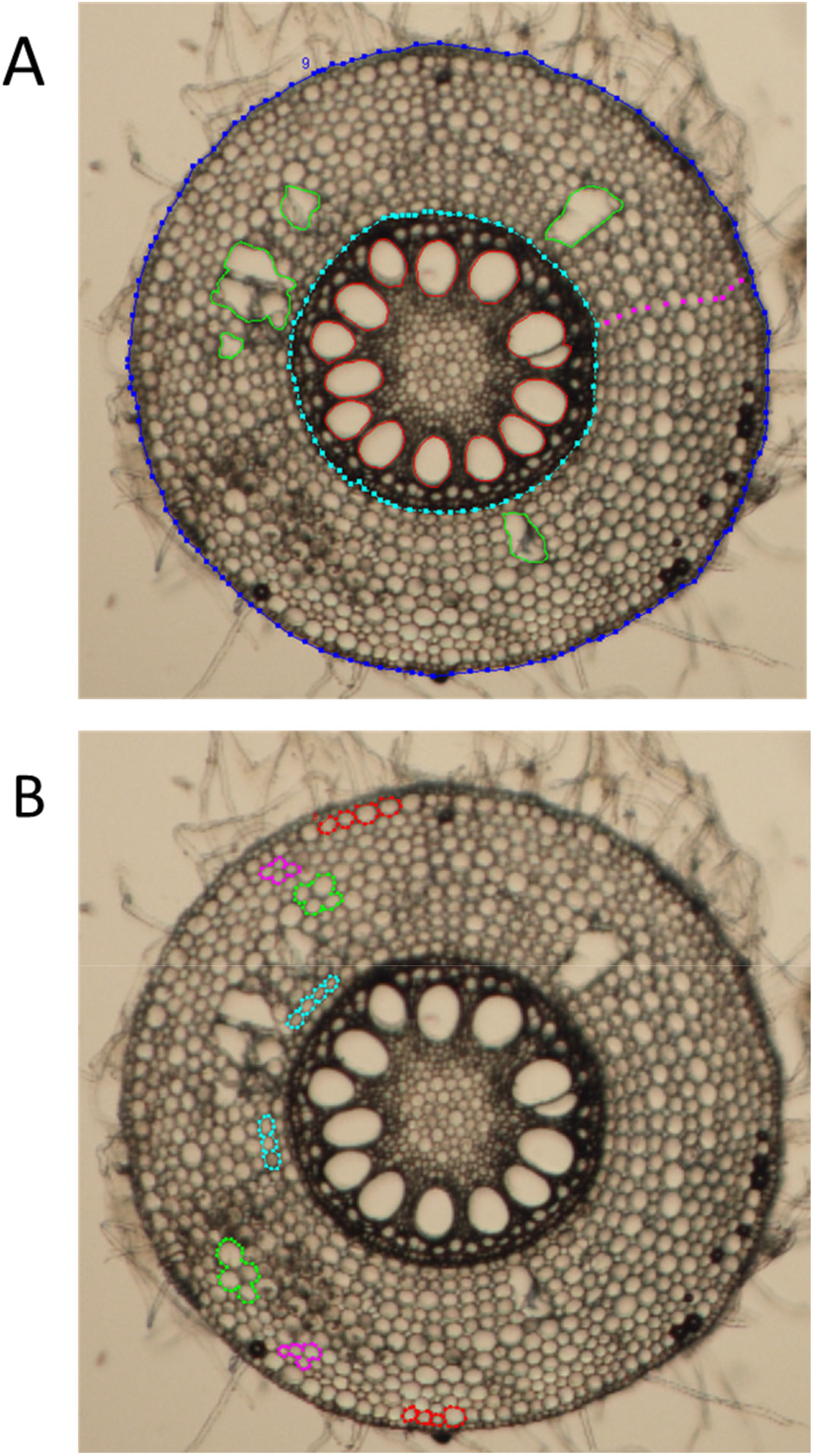
Maize root cross-section image analysis. An ObjectJ macro in ImageJ was created to semi-automate analysis of root-cross section images. The same anatomical traits were analyzed in LAT and manually sectioned roots, with minor differences in methodology. An example of an analyzed image from manual cross-sections. (A) The root (dark blue, outer), stele (cyan, inner), and aerenchyma (green) were outlined and total areas and ratios calculated. Individual metaxylem vessels were traced (red) and individual and total areas determined. Vessels beginning to divide were traced as a single vessel. Images were zoomed in to allow accurate tracing along the outer edge of each vessel. Cortical cell file number was manually counted in three positions to account for root asymmetry, and a representative axis across the cortex was selected to record a representative cell file count (the count of pink points) and measure cell diameters of each cell file (distance between every consecutive pink point). The innermost cell layer (i.e. distance between the innermost pink point and the stele boundary) was often incomplete and was not recorded. The cell diameters were used to calculate the cell sizes of the hypodermis (HYP), outermost (OUT) and innermost (INN). (B) For one full replicate, the inner (cyan), mid (green), outer (pink), and hypodermis (red) cortical cell sizes were determined and used to validate estimates from cell diameters as described above. Up to eight representative cells per layer were traced and average cell cross-sectional areas were calculated. Images were zoomed in to allow careful tracing of the outer edge of the cells; cell walls were included in the trace. Unlike LAT images, manual cross-sections from greenhouse-grown plants showed intact epidermis cells and root hairs.

**Fig S3.**
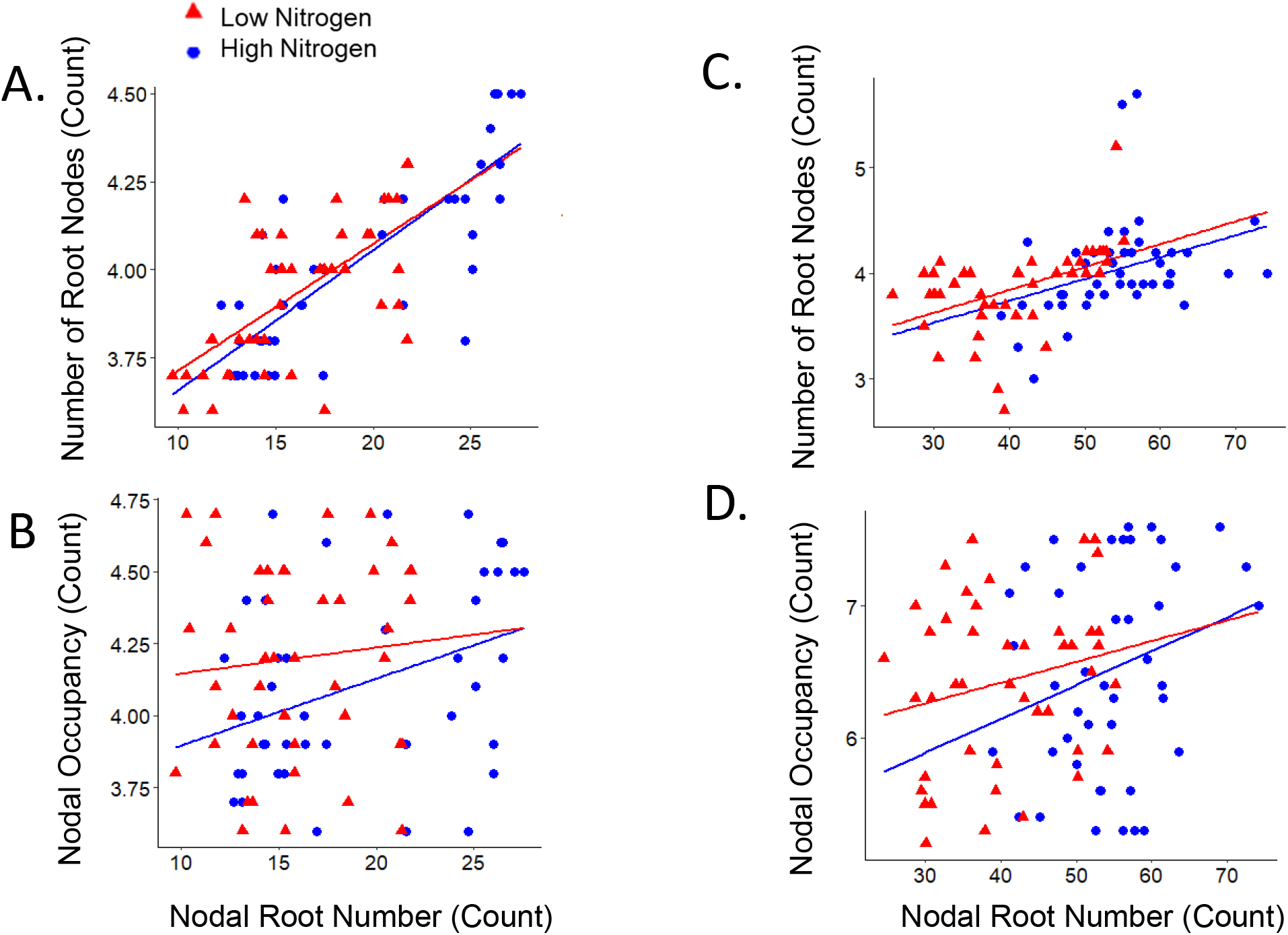
Genotypic contrast in nodal root number and diameter among maize RILs in the field and greenhouse. (A) total number of nodal roots emerged at harvest and number of root nodes, (B) nodal root number and nodal occupancy from nodes 2, 3, and 4 in the (C) total number of nodal roots emerged at harvest and number of root nodes, (D) nodal root number and nodal occupancy from nodes 1, 2, and 3 in the field. High and low nitrogen treatments are indicated (HN, blue; LN, red).

**Figure S4.**
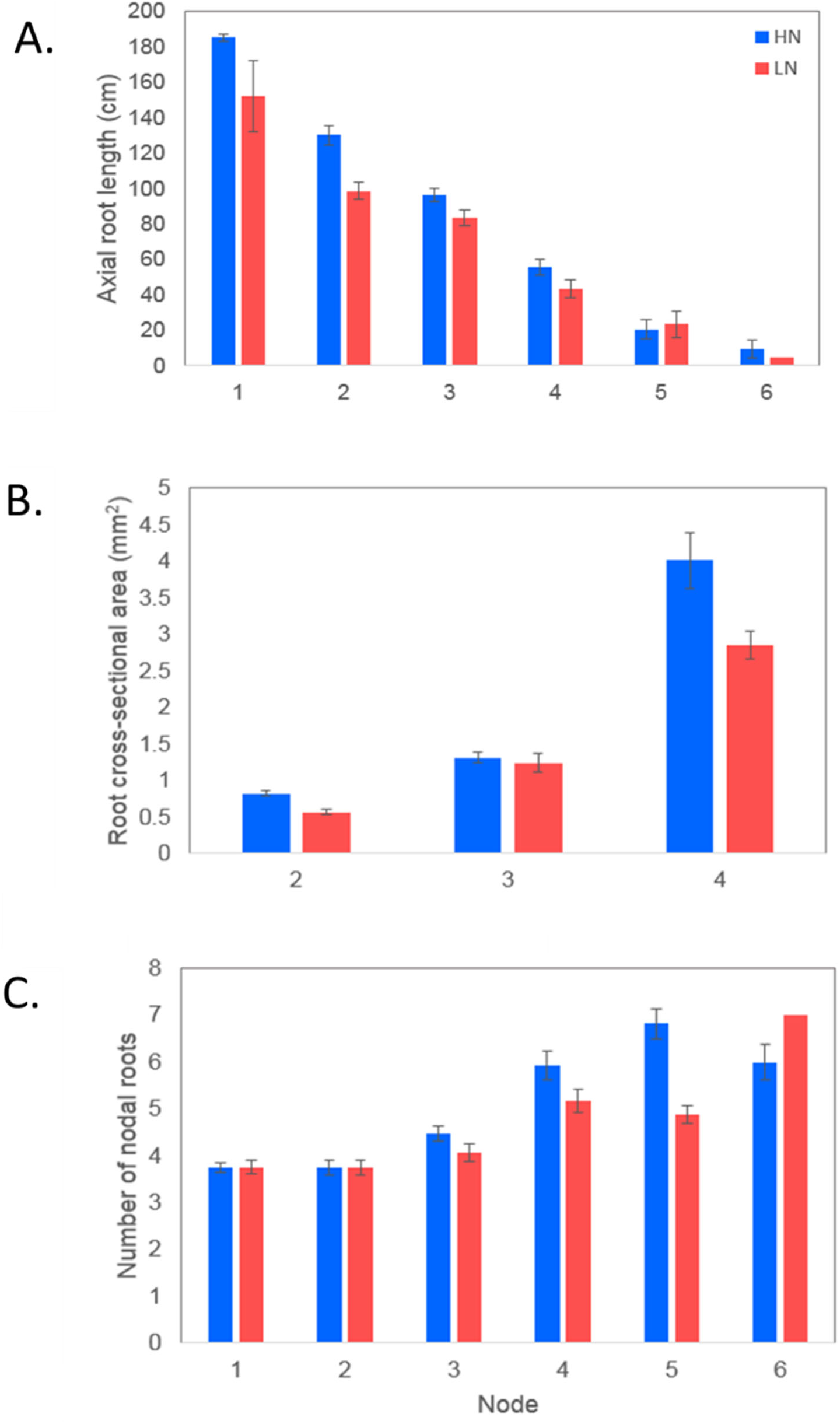
Axial root lengths, diameters, and occupancy by node under high and low nitrogen in IBM RILs. Means ± SE of (A) the average axial root length (ARL) by node at time of harvest, and (B) average RXA by node, and (C) average number of roots per node (NO), in the greenhouse. ARL was evaluated in all plants for nodes 2 through 6, and only a subset of plants in node 1. High and low nitrogen treatments are indicated (HN, blue; LN, red).

**Figure S5.**
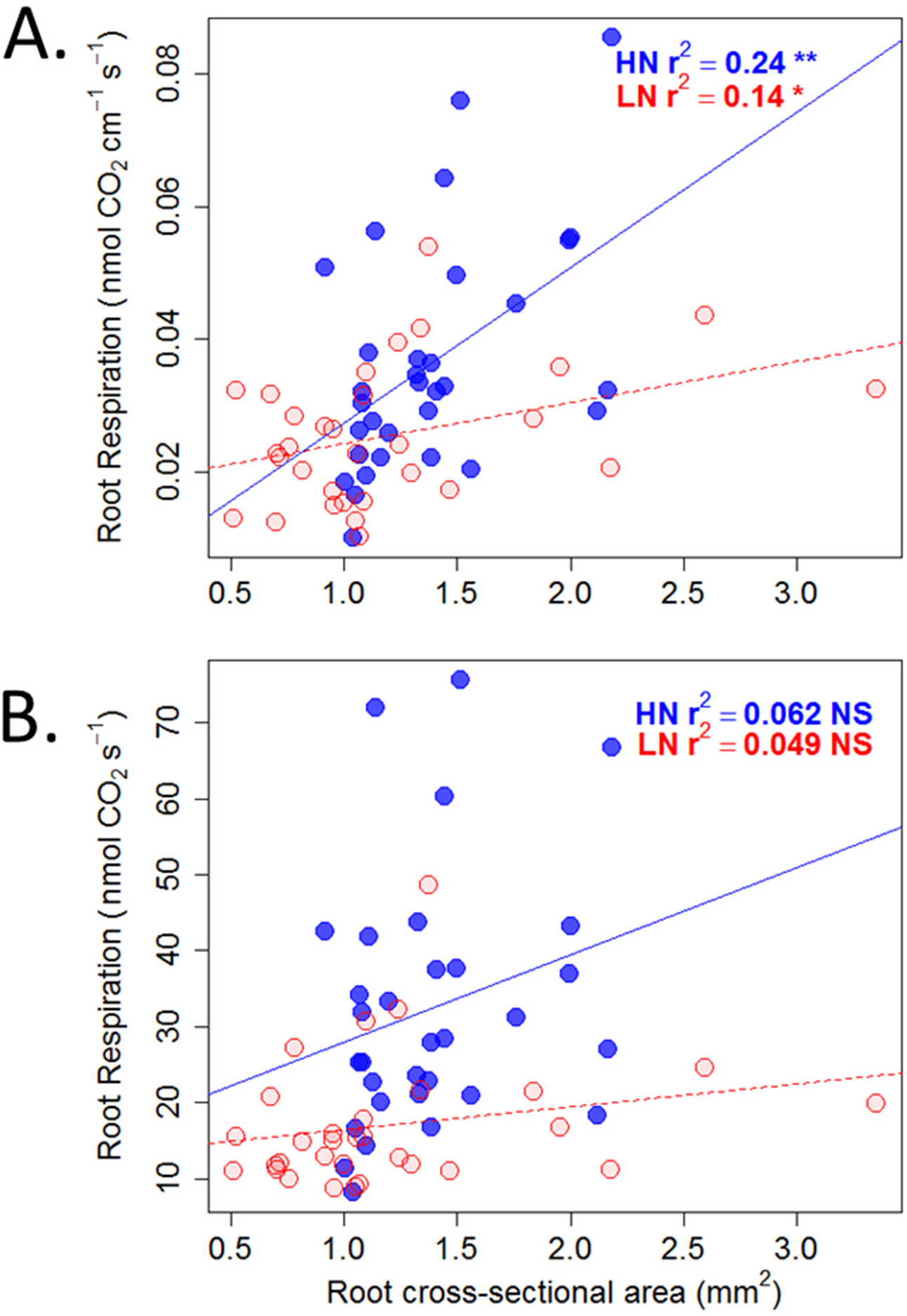
Relationship between axial root diameter and respiration among maize RILs. Linear regression of root cross-sectional area (RXA) averaged from second and third node roots against (A) root respiration per unit root length averaged from three roots each from nodes 2 and 3, and (B) total axial root respiration (root respiration rate multiplied by axial lengths of all developed nodes except the first node), from individual plants of maize IBM RILs grown in high (HN, blue) or moderate low nitrogen (LN, red) treatments in the greenhouse. R^2^ value and significance (p< 0.05*, 0.01**, p>0.1 NS, not significant) are indicated.

